# Integrating gene drives with established pest controls to mitigate spillover risk

**DOI:** 10.1101/2025.09.15.676306

**Authors:** Sviatoslav R. Rybnikov, Adam Lampert, Gili Greenbaum

## Abstract

Gene drives—artificial genetic constructs that can rapidly spread in a population due to biased inheritance—are a promising tool for controlling disease vectors and invasive species. While gene drives can eradicate target populations within several generations, their unintended spillover into non-target populations poses serious ecological concerns. Here, we examine how integrating gene drives with established pest control methods, such as pesticide application and sterile-male release, can mitigate spillover risk. We develop optimal control models to identify cost-effective treatment strategies that minimize total eradication costs, accounting for actual investments and potential damage from spillover. We show that pesticides are most cost-effective at the intermediate stage of eradication when gene drives have already slowed population growth but do not yet substantially suppress the population. For sterile-male release, however, we observe two distinct types of optimal strategies, the choice of which depends on parameter values: intensive early-stage releases, or mild later-stage releases. Using established pest controls also relaxes requirements to gene drive design, favoring milder suppressive effects as the treatment intensifies.

## 1. INTRODUCTION

Pest populations such as disease-vector species, crop-damaging species, and invasive species cause significant ecological and economic damage. Over the last century, several pest control methods have been developed, including chemical pesticides, the release of sterile males, and biological control methods. Another approach being currently developed involves the use of gene drives—genetic elements that violate standard Mendelian inheritance laws (Alphey *et al*. 2020) and can spread rapidly in populations (Burt 2003; Deredec *et al*. 2008; Unckless *et al*. 2015). Since gene drives could induce substantial fitness costs to the individuals carrying them, they are being explored as a promising tool to suppress harmful populations. Gene drives are currently being developed for weeds (Barrett *et al*. 2019; Liu *et al*. 2024; Oberhofer *et al*. 2024) and rodents (Gierus *et al*. 2022; Manser *et al*. 2019; Pfitzner *et al*. 2020), and have reached advanced stages of laboratory experiments in mosquitoes (Hammond *et al*. 2021; Hammond & Galizi 2017; Nolan 2021). However, despite its potential, gene drive technology raises substantial concerns (Esvelt & Gemmell 2017; Frieß *et al*. 2023; Hartley *et al*. 2022; Jiang & Wang 2024; Schwartz *et al*. 2025), such as the risk of unintended spillover of gene drive alleles to non-target populations, which might severely damage ecosystems (Kim *et al*. 2023; Noble *et al*. 2018; Oye *et al*. 2014; Webber *et al*. 2015). Therefore, approaches for mitigating spillover risk must be developed for gene drive technology to move toward field application.

The dynamics of populations carrying gene drives have been modeled in numerous theoretical studies (reviewed in Frieß *et al*. 2023; Kim *et al*. 2023) that have focused on conditions under which spillover occurs, and explored how to mitigate it. For example, several studies have identified ranges of gene drive characteristics (fitness cost, conversion rate, etc.) that result in threshold-dependent spread, when the drive can invade only if introduced above a certain frequency, reducing risk of unintentional introductions (Burt 2003; Deredec *et al*. 2008; Leftwich *et al*. 2018; Unckless *et al*. 2015). Other studies have explored sophisticated genetic architectures that may allow gene drive confinement to limited geographic localities (Champer *et al*. 2020; Noble *et al*. 2019; Zhu & Champer 2023), or how gene drive sensitivity to environmental conditions can be leveraged to mitigate spillover risk (Girardin *et al*. 2019; Harris & Greenbaum 2023; Marshall & Akbari 2018; Tanaka *et al*. 2017). Recent studies have coupled evolutionary (allele-frequency) dynamics with demographic (population-size) dynamics (Beaghton & Burt 2022; Girardin & Débarre 2021; Harris & Greenbaum 2023; Kläy *et al*. 2023). This is particularly important for gene drives aimed at suppressing populations, as the interplay between demographic and evolutionary processes may strongly affect spillovers. The inclusion of demography in gene drive models also opens the possibility for modeling how other processes that affect population sizes, such as traditional control measures, can interact with the dynamics of gene drives.

While most previous studies mainly considered gene drive as the only control measure applied, it is often advantageous to combine several control measures together. Combining several controls is a common practice in established pest control, where treatment schemes involving pesticide application, sterile-male release, and pheromone-based mating disruption have repeatedly proven effective in the field (Bansal *et al*. 2024; Gutierrez *et al*. 2019; Horner *et al*. 2020; Kittayapong *et al*. 2019), and theory has been developed for various such combinations (Barclay 1980; Lampert & Liebhold 2021; Larramendy & Soloneski 2012; Tang & Cheke 2008). Therefore, integrating gene drives with established control measures may open avenues to leverage their interactions, maximizing advantages and improving the cost-effectiveness of population suppression.

Here, we examine the potential of complementing gene drives with two key established pest controls: pesticide application and sterile-male release. Our goal is to identify optimal treatment strategies—the timing and intensity of different interventions that minimize the total cost of eradication over time, considering both the cost of direct expenditures and the cost due to spillover risk. Specifically, we consider the risk of spillover to be proportional to the overall number of gene drive carriers through the entire eradication process. While such cost minimization, known as bioeconomic modelling, has become a common approach for established pest controls, it remains almost overlooked in gene drive models, with only a few simulation studies incorporating cost (e.g., assessing health care outcomes related to insecticide-treated nets and vaccinations in (Hancock *et al*. 2024; Metchanun *et al*. 2022)). In addition to identifying optimal treatment strategies for given gene drives, we examine how these strategies depend on biological and economic parameters, including drive configurations and population growth rates.

## 2. MODEL AND METHODS

### 2.1. Demographic gene drive model

To study how to combine multiple control methods optimally, we developed a model that incorporates both evolutionary dynamics (i.e., changes in the frequency of the gene drive allele) and demographic dynamics (i.e., changes in the population density) of a target population. As a baseline for evolutionary dynamics, we use a model of a standard homing gene drive with gametic conversion that is expected to reach fixation when released in a population (Deredec *et al*. 2008; Unckless *et al*. 2015). As a baseline for the demographic dynamics, we use a standard discrete-time model of population growth under intraspecific competition (Beverton & Holt 1957). We first describe the dynamics of a population in which a gene drive is deployed without additional controls, and in the following subsection we describe how additional pest controls, such as pesticides and sterile males, modify the dynamics.

We consider a random-mating diploid population with non-overlapping generations. Spillover of the gene drive to non-target populations might occur, but we do not model other populations explicitly. We consider a single locus with two alleles: wild-type allele and gene drive allele. We assume that the gene drive allele reduces the fitness of its carriers with relative fitness cost *s* and dominance *h*, such that gene drive homozygotes have fitness 1–*s*, heterozygotes have fitness 1–*hs*, and wild-types have fitness 1. To model the gene drive mechanism, we assume that heterozygotes are converted to gene drive homozygotes with probability c; we assume that conversion occurs during gamete formation, in accordance with current empirical data (Grunwald *et al*. 2019; Kyrou *et al*. 2018). Under these conditions, the frequency of the gene drive allele, *q*_*t*_, can be formulated as (Deredec *et al*. 2008; Unckless *et al*. 2015):

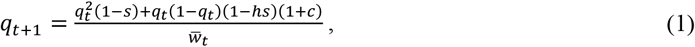

where the numerator sums the contribution of the genotypes in terms of gene drive alleles, and the denominator represents the mean population fitness, 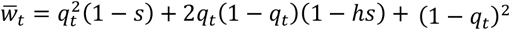. We show in Appendix A that Eq. 1, which was previously derived by considering gene drive frequency alone, also applies to the setup explicitly considering both gene drive frequency and population density. Depending on the gene drive configuration (*s, c, h*), these dynamics can result in several outcomes: fixation of the gene drive, loss of the gene drive, a stable equilibrium with both gene drive and wild type alleles present, or bistability with either fixation or loss of the gene drive – depending on initial conditions (Deredec *et al*. 2008; Unckless *et al*. 2015). Here, we focus on configurations that result in fixation of the gene drive allele regardless of its initial frequency.

To model the impact of gene drives on the population density, we adjust a standard population growth model to account for the reduced fitness of the gene drive carriers. Specifically, we assume that, in the absence of gene drive carriers, the population grows according to the Beverton-Holt model (Beverton & Holt 1957), which is a standard discrete-time analog of the continuous logistic model:

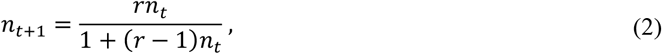

where, *n*_*t*_ is the population density at time *t* (relative to the carrying capacity), and *r* is the intrinsic growth rate without intraspecific competition of a population of wild-type individuals. In turn, as the proportion of the gene drive allele in the population increases, fewer individuals survive and reproduce; therefore, we model the population growth rate to be proportional to the mean population fitness, and modify the Beverton-Holt model accordingly (full derivation in Appendix A):

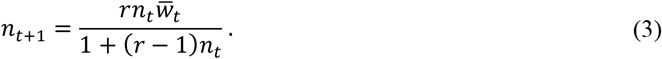

For a wild-type population, which has a mean fitness of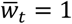, Eq. 3 reduces to Eq. 2. In the presence of the gene drive, 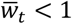, and the population grows slower and may even decline.

### 2.2. Dynamics under additional pest controls

The dynamics in Eq. 3 model the population size under the impact of the gene drive spread, without any additional interventions. Next, we extend the model by incorporating established pest controls: pesticides and sterile males. For pesticides, we assume that a certain amount of chemical pesticides removes a fixed fraction of the population (Clark 2010; Lampert & Liebhold 2021). As a scaled unit, we consider a single pesticide application to be the amount that reduces the population size by half. We denote the number of pesticide application units used at time *t* as *P*_*t*_ (*P*_*t*_ ≥ 0), and we adjust Eq. 3 to incorporate pesticide application:

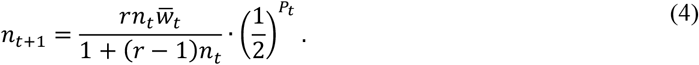

Next, we model the effect of sterile-male releases on the dynamics. We denote the amount of sterile males released at time *t* as *M*_*t*_ (*M*_*t*_ ≥ 0), measured in units of fractions of the carrying capacity to be compatible with the population density *n*_*t*_. Assuming a 1:1 sex ratio and that each female mates once, the proportion of females encountering a fertile mate is 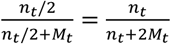 (Knipling 1959; Lampert & Liebhold 2021; Sawyer *et al*. 1987). We can adjust the numerator in Eq. 4 by this proportion to formulate the population size after sterile-male release:

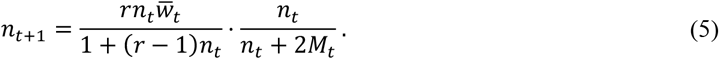

Finally, when both pesticides and sterile males are applied to a population with a gene drive, the joint effect of the three control measures can be formulated as:

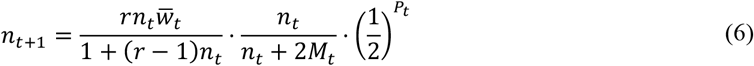

### 2.3. Formulating spillover risk mitigation as an optimal control problem

To examine how to optimally combine gene drives with the established pest control methods, we formulate an optimal control problem. The need for optimization arises because using established pest controls bears a trade-off: they speed up eradication and thereby reduce the number of potential gene drive carriers escaping to non-target populations (Fig. 1) but inevitably increase the cost of eradication.

**Figure 1.**
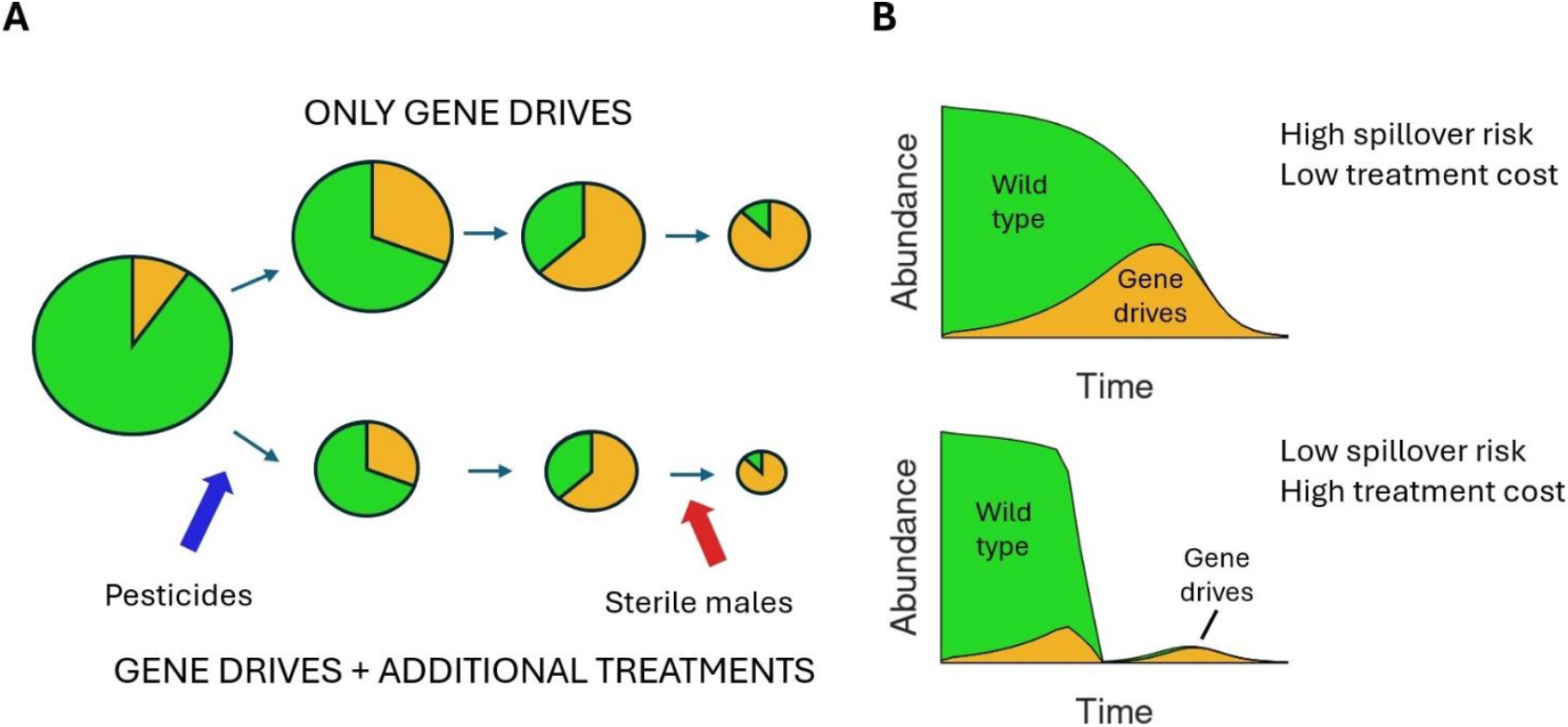
Illustration of the gene drive dynamics, with and without additional treatment methods. (A) Circles represent the population at different times throughout the treatment process (from left to right) or following different treatment scenarios (upper row versus bottom row). Circle sizes represent population size, and their colors represent the genetic composition (orange is gene drives, green is wild-type). In both top and bottom rows, gene drives spread over time (the orange area becomes larger relative to the green area), and the population shrinks until it vanishes (circle sizes decrease over time). The top row demonstrates the dynamics without additional treatment (no pesticides and no sterile males). In the bottom row, pesticides and sterile males are introduced at different stages to speed-up the eradication process. (B) Demonstrated are the abundances of the gene drive carriers and the wild-type individuals during eradication, without (top row) and with (bottom row) additional treatments. Spillover risk is proportional to the total number of gene drive carriers over time, represented approximately by the orange area in the population dynamics plots. Adding established pest controls (e.g., pesticides, sterile males) accelerates population suppression and thereby reduces spillover risk compared to using gene drives alone (top row vs. bottom row). However, these additional interventions incur their own costs, and minimizing the total cost can be optimized by controlling the timing and intensity of treatment application.

Our objective is, therefore, to minimize the *net present cost* (NPC)—the total cost of eradication over time, which includes (i) the cost due to spillover risk, (ii) the amount invested in pesticides, and (iii) the amount invested in sterile males. To quantify the cost due to spillover, we assume that each gene drive carrier has a fixed probability of escaping to non-target populations. Accordingly, the spillover risk in a given generation *t* is proportional to the number of gene drive carriers in that generation, *n*_*D,t*_, and the total spillover risk is proportional to the total number of gene drive carriers over the entire process (Fig. 1). Specifically, we equally account for all gene drive carriers, both homozygotes and heterozygotes, and use the Hardy-Weinberg genotype frequencies:

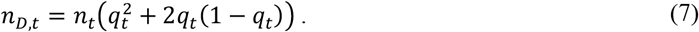

In turn, the amount invested in additional pest controls is proportional to pesticide application in each generation, *P*_*t*_, and sterile-male release in each generation, *M*_*t*_. Thus, NPC is given by:

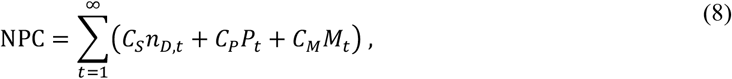

where *C*_*S*_ is the per-unit cost due to potential spillover, *C*_*P*_ is the per-unit cost of pesticide application, and *C*_*M*_ is the per-unit cost of sterile-male release. Higher *C*_*S*_ values could result from higher migration rates to non-target populations, a higher probability of spillover occurring if a gene drive carrier migrates to a non-target population, and a more substantial evaluation of the damages of a potential spillover. Note that we do not consider discounting in Eq. 8, since eradication takes only several weeks, and the difference between present and delayed expenditures is negligible.

We scale the cost coefficient to the risk of spillover and consider only two cost ratios: *α* = *C*_*P*_/*C*_*S*_ and *β* =*C*_*M*_/*C*_*S*_; hereafter, these relative costs are referred to as the *per-unit treatment costs*. Higher *α* and *β* values imply higher costs of pesticides and sterile males, respectively, relative to the cost due to spillover risk. Our optimization procedure is conducted through dynamic programming, described in detail in Appendix B. Since the per-unit cost due to potential spillover is largely unknown and hard to estimate, we consider a wide range of values of *C*_*S*_ (or *α* and *β*). This enables us to find the minimal total spillover risk that can be achieved as a function of the total investment in the other control methods. This multiple-objective optimization approach enables the examination of the full range of possibilities – not only for a certain choice of the relative cost.

### 2.4. Model parameterization

We focus on ‘successful’ suppression gene drives, which invade the target population regardless of their initial frequency and eventually eradicate it. For tractability, most shown results are for a gene drive with configuration (*s, c, h*) = (0.7, 0.9, 0.5) deployed in a population with intrinsic growth rate *r* = 2. This baseline gene drive configuration is a reasonable representation for highly efficient suppression gene drives currently being tested in controlled laboratory conditions (Gantz *et al*. 2015; Hammond *et al*. 2016; Kyrou *et al*. 2018). We also examine deviations from the chosen baseline setting by varying fitness cost *s* of the gene drive (ranging from 0.3 to 1) and intrinsic growth rate *r* of the population (ranging from 1.1 to 10). When varying these parameters, we only consider settings that result in gene drive spread and population eradication. For example, with an intrinsic growth rate of *r* = 2, we consider the fitness cost range from *s* = 0.5 (below which the population is not suppressed by the gene drive) to *s* = 0.95 (above which gene drives fail to spread). We mainly report results for a standard deployment scenario, where a small number of gene drive homozygotes (*q*_0_ = 0.01) are introduced into a large population stable at the carrying capacity (*n*_0_ = 1); however, to understand which population states require additional treatments, we also explore the whole space of possible initial conditions, with both *q*_0_ and *n*_0_ ranging from 0.01 to 1. Since it is unclear how spillover damage should be quantified, we examine a range of scenarios, from the cost of spillover being negligible relative to that of treatment (*α, β* → ∞) to it being immeasurably large (*α, β* = 0).

## 3. RESULTS

### 3.1. Pesticide applications are most cost-efficient during peak gene drive abundance

First, we investigate the optimal strategies for combining gene drives with pesticide application. Our results show that, if the per-unit treatment cost *α* is below a certain threshold (i.e., treatment is relatively cheap), the optimal strategy is to rapidly lower the population size with pesticides (*α =* 0 in Fig. 2A). Above this threshold, however, the optimal treatment strategy utilizes both gene drives and pesticides, with pesticide application beginning only a few generations after the gene drive release (Fig. 2A). This lag allows gene drives to increase in frequency and reduce the population’s growth rate. As *α* increases further, the optimal strategies utilize pesticide applications only later on and become milder in their intensity. Finally, if *α* is above an even higher threshold, pesticide applications are no longer cost-effective, and the optimal strategy is to allow gene drives to lead to eradication without further intervention.

**Figure 2.**
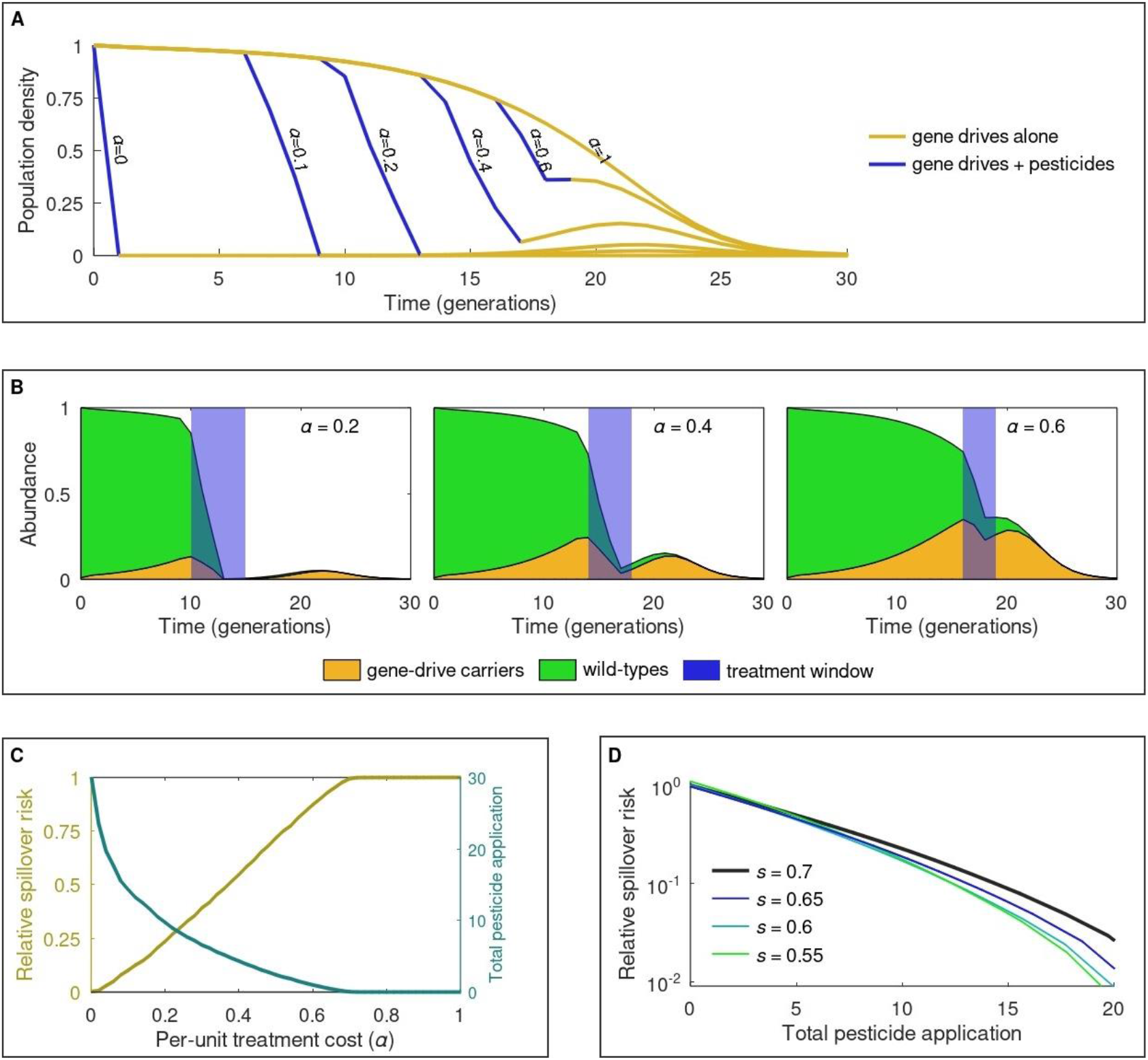
Optimal strategies for minimizing spillover risk by combining gene drives with pesticides. (A) Demonstrated are population dynamics under optimal pesticide application for different per-unit treatment costs (*α*), where the dynamics follow Eqs. 1 and 4. Each curve corresponds to a certain value of *α* denoted next to it. (B) Demonstrated are genotype dynamics and treatment time windows for the optimal strategies for three values of *α*. (C) Relative spillover risk and total pesticide application following the optimal solution as functions of *α*. (D) Spillover-treatment curves, showing the minimal spillover risk achievable for a given amount of available pesticide, for different values of *s*. Spillover risk shown on the y-axis is relative to the spillover risk of the baseline gene drive (*s* = 0.7) under no pesticide treatment. The curves are obtained by identifying the optimal treatment strategy for numerous values of *α* and computing relative spillover risk against total pesticide application. Parameters: *c* = 0.9, *h* = 0.5, *r* = 2; in (A-C) *s* = 0.7. Initial conditions are *n*_0_ = 1 and *q*_0_ = 0.01.

These results imply that both very early and very late pesticide applications are generally less cost-effective. At early stages, when gene drive carriers are rare, an initially high growth rate requires repeated pesticide applications and incurs additional costs. At late stages, when low fitness gene drive carriers are abundant, the gene-drive-induced negative growth rate ensures eradication without pesticides. In addition, pesticide applications are avoided at the early and late stages because of the low abundance of gene drive carriers, and hence low spillover risk, at these stages (Fig. 1B). Thus, pesticide applications are cost-effective when the number of gene drive carriers (and hence the spillover risk) reaches its peak under a no treatment scenario, i.e. when the gene drive allele has already spread significantly but has not yet suppressed the population substantially. As pesticides become cheaper (lower *α* values), the treatment time window broadens (Fig. 2B).

Following the optimal solution, as the per-unit treatment cost increases, the total amount of pesticide application decreases, while the spillover risk increases (Fig. 2C). Specifically, the curve showing the resulting spillover as a function of the total pesticide application (Fig. 2D) represents a Pareto front, in which one cannot obtain a solution that both requires fewer pesticide applications and results in lower spillover. The obtained spillover-treatment curve is convex, suggesting diminishing marginal efficiency from increasing pesticide application (Fig. 2D), where each value of α, results in a particular choice of total investment along the Pareto front.

Notably, cost-effectiveness can further be improved if the gene drive configuration is also subject to optimization. A key parameter that substantially affects gene drive dynamics is its fitness cost, *s* (Deredec *et al*. 2008; Unckless *et al*. 2015). Since gene drives with higher *s* values spread more slowly but suppress the population more strongly, an optimal fitness cost *s*\* must exist that minimizes the number of gene drive carriers over eradication, and hence also the spillover risk. Comparing spillover-treatment curves for various *s* values shows that, when the total pesticide application is zero, the optimal fitness cost is *s*\* ≈ 0.65, slightly below the baseline configuration (*s* = 0.7). As intervention intensity increases, the optimal fitness cost tends to decrease (Fig. 2D). Thus, combining gene drives with pesticide application favors gene drives with lower fitness costs, which could provide additional risk mitigation, especially under moderate-to-strong interventions.

### 3.2. Sterile-male strategies shift abruptly from weak to strong treatment regimes

Next, we consider the combination of gene drives with sterile-male release. Similarly to pesticides, for higher per-unit treatment cost *β*, the optimal strategy results in smaller total sterile-male release and higher relative spillover risk (Fig. 3). However, we observe two qualitatively different types of optimal strategies which shift abruptly as *β* surpasses a critical threshold *β*_*c*_ – a behavior not observed for pesticides. For *β* < *β*_*c*_, the optimal strategy is to release a large number of sterile males already during the first few generations following the gene drive release (Fig. 3A, trajectories on the left). For higher *β* values (closer to the threshold), the releases start later and stop earlier, resulting in fewer sterile males being released overall. For *β* > *β*_*c*_, the optimal strategy changes to minimal, if any, release of sterile males at a later stage, preceding the final stage where gene drive completes eradication alone (Fig. 3A, trajectories on the right).

**Figure 3.**
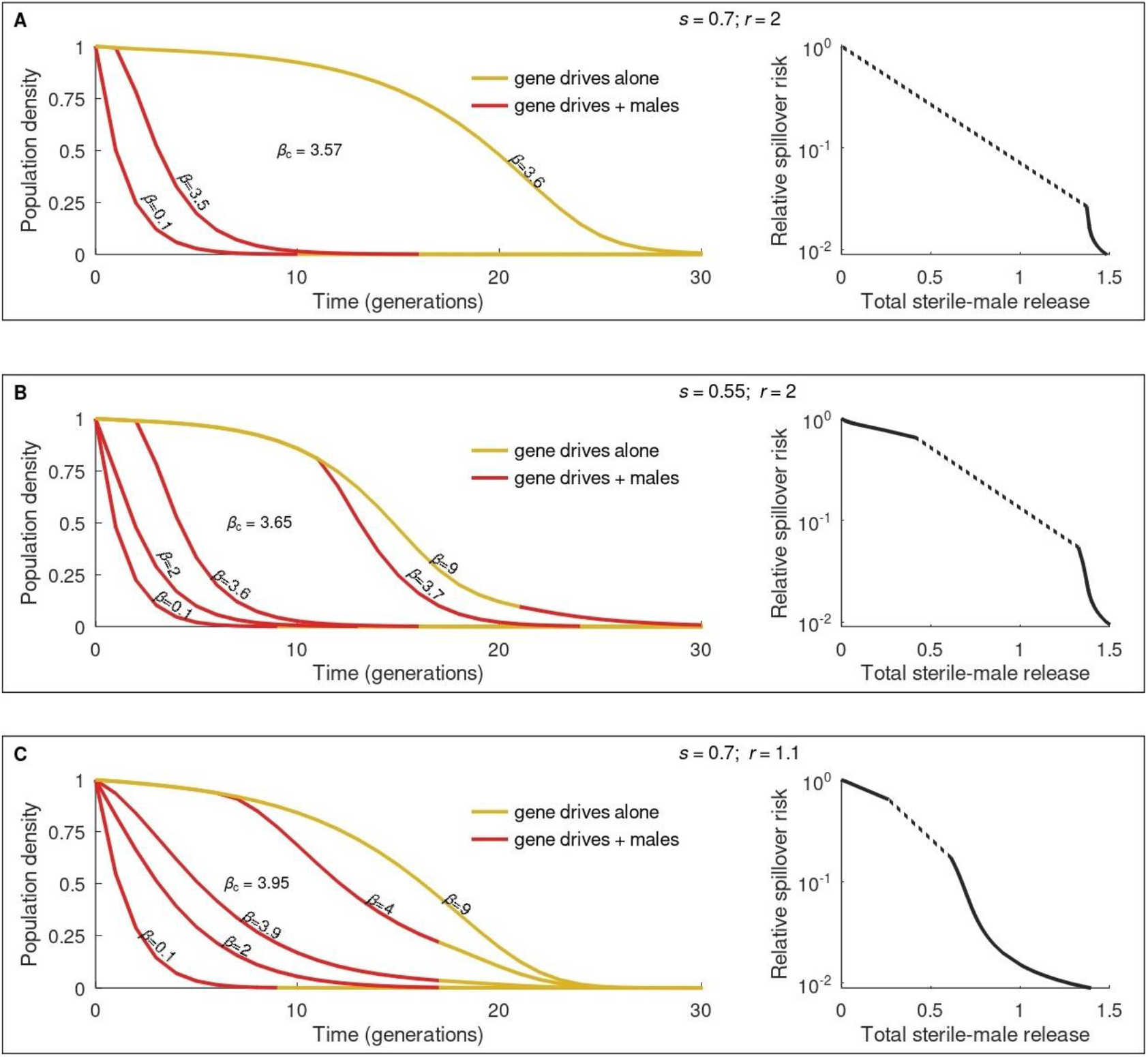
Optimal strategies for minimizing spillover risk by combining gene drives with sterile-male releases. Each box (A-C) shows optimal solutions for a different choice of the parameters *r* and *s*. In each box, the left panel shows the population dynamics under optimal sterile-male release for different per-unit treatment costs (*β*), denoted next to each curve, where the dynamics follow Eqs. 1 and 5. At a critical value *β*_c_, the optimal treatment strategies change abruptly (the value of *β*_c_ is noted between the relevant curves). The right panels show the spillover-treatment curve, showing the minimal spillover risk that can be achieved for a given amount of sterile males available. The dashed segments show ranges of sterile-male release that cannot be chosen as the optimal solution. Parameters: *c* = 0.9 and *h* = 0.5. Initial conditions: *n*_0_ = 1 and *q*_0_ = 0.01.

Not only do treatment strategies shift abruptly at *β* = *β*_*c*_, but also the two outcomes following the optimal solution: total sterile-male release and relative spillover risk. The abrupt transition leaves a region of the spillover–treatment curve uncovered because optimal strategies with intermediate sterile-male release do not emerge (Fig. 3, dashed segments in the right panels). This implies that the curve is concave in that region, and therefore, there is no value of the marginal cost *β* for which it is optimal to choose treatment from that region. The range of unobserved releases depends on parameter values; for example, it is narrower under lower fitness costs (Fig. 3B) and under lower intrinsic growth rates (Fig. 3C).

The observed qualitative difference between pesticides and sterile males—gradual vs. abrupt changes in optimal strategies—can be considered from the perspective of population states requiring treatment. To be cost-efficient, pesticides require intermediate gene drive frequencies and sufficiently high population densities, and these conditions become stricter as the per-unit treatment cost increases. The interplay between population dynamics and intervention efficacy can be visualized on a phase plane, where axes represent the abundances of gene drive and wild-type alleles. On such a plane, the above-mentioned conditions outline a triangular area adjacent to the hypotenuse; the area progressively contracts with *α*, gradually reducing the proportion of the population’s trajectory that intersects it (Fig. 4A, blue). In contrast, sterile males are cost-efficient when both gene drive frequency and population density are low enough. These conditions contour a drop-shaped area adjacent to the left edge of the phase plane. The area moves downward with *β*, so at a critical *β* value, the entire trajectory shifts from lying inside (Fig. 4B, left) to lying mostly outside (Fig. 4B, right).

**Figure 4.**
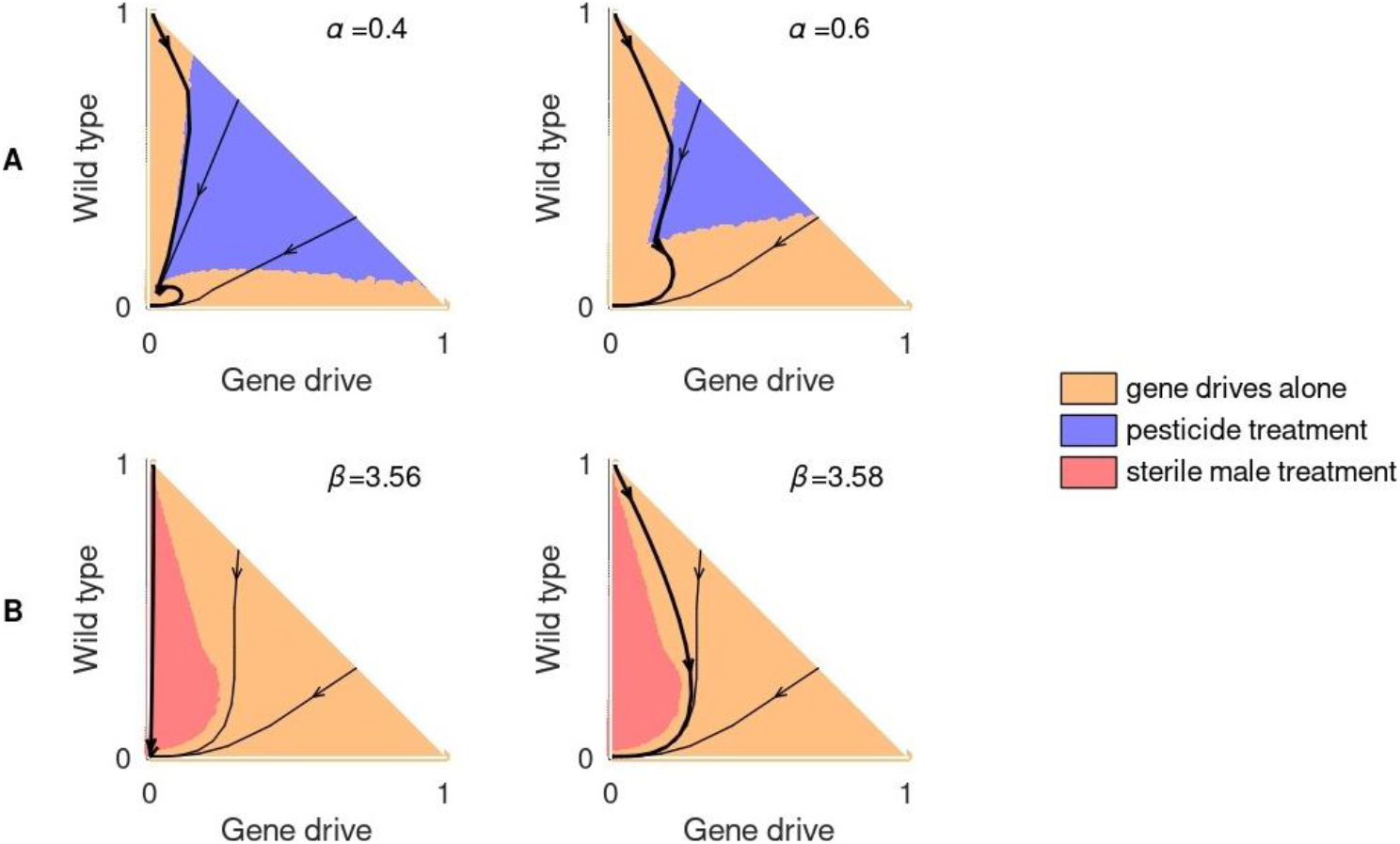
Population phase diagrams for gene drives combined with another treatment method. (A) Combining gene drives with pesticide application (Eq. 4). (B) Combining gene drives with sterile-male release (Eq. 5). The background colors denote states where gene drives are used alone (orange) or combined with pesticides (blue) or sterile males (red). For the shown gene drive configuration in (B), the critical per-unit treatment cost of sterile males is *β*_c_ ≈ 3.57 (as in Fig. 3A), and the two phase diagrams show scenarios slightly below (left) and slightly above (right) the threshold. Axes represent abundances of the two alleles relative to the carrying capacity. The black curves show the population’s trajectories starting at different initial conditions (the bold curve corresponds to the initial conditions considered throughout most of the paper). Parameters: *c* = 0.9, *s* = 0.7, *h* = 0.5, and *r* = 2.

### 3.3. For combined optimal treatments, pesticides are used early and sterile males later

Finally, we study the more general scenario, where both pesticides and sterile males can be used to mitigate spillover risk. Because we are now considering two possible interventions, we consider both per-unit treatment costs: α for pesticide application and *β* for sterile-male release. The ratio between these two parameters, *γ* = *α*/*β*, determines whether a single treatment method or their combination is favored. Specifically, we observe two thresholds, *γ*_1_ and *γ*_2_ (*γ*_1_ ≤ *γ*_2_), where pesticides are cost-effective when the ratio *γ* is below the upper threshold (*γ* ≤ *γ*_2_) and sterile males when *γ* is above the lower threshold (*γ* ≥ *γ*_1_). Thus, both treatment methods should be applied only in the intermediate range *γ*_1_ ≤ *γ* ≤ *γ*_2_ (see Eq. B4 in Appendix B).

We found that when the per-unit treatment cost of one control method is low (relative to the other method and spillover risk), only this method complements gene drives if needed. This scenario is demonstrated in the top row and left column of Fig. 5: sterile males are never used when *α* = 0.01, whereas pesticides are never used when *β* = 0.1 (excluding the top-left panel, where *α* and *β* differ the least). Conversely, when the per-unit treatment cost of a method is substantially higher than that of the other method, it is abandoned, and the resulting pattern mirrors using the cheaper method alone. For example, the right column in Fig. 5 resembles the pesticides-only scheme (Fig.4A), while the bottom row aligns with the males-only scheme (Fig. 4B).

**Figure 5.**
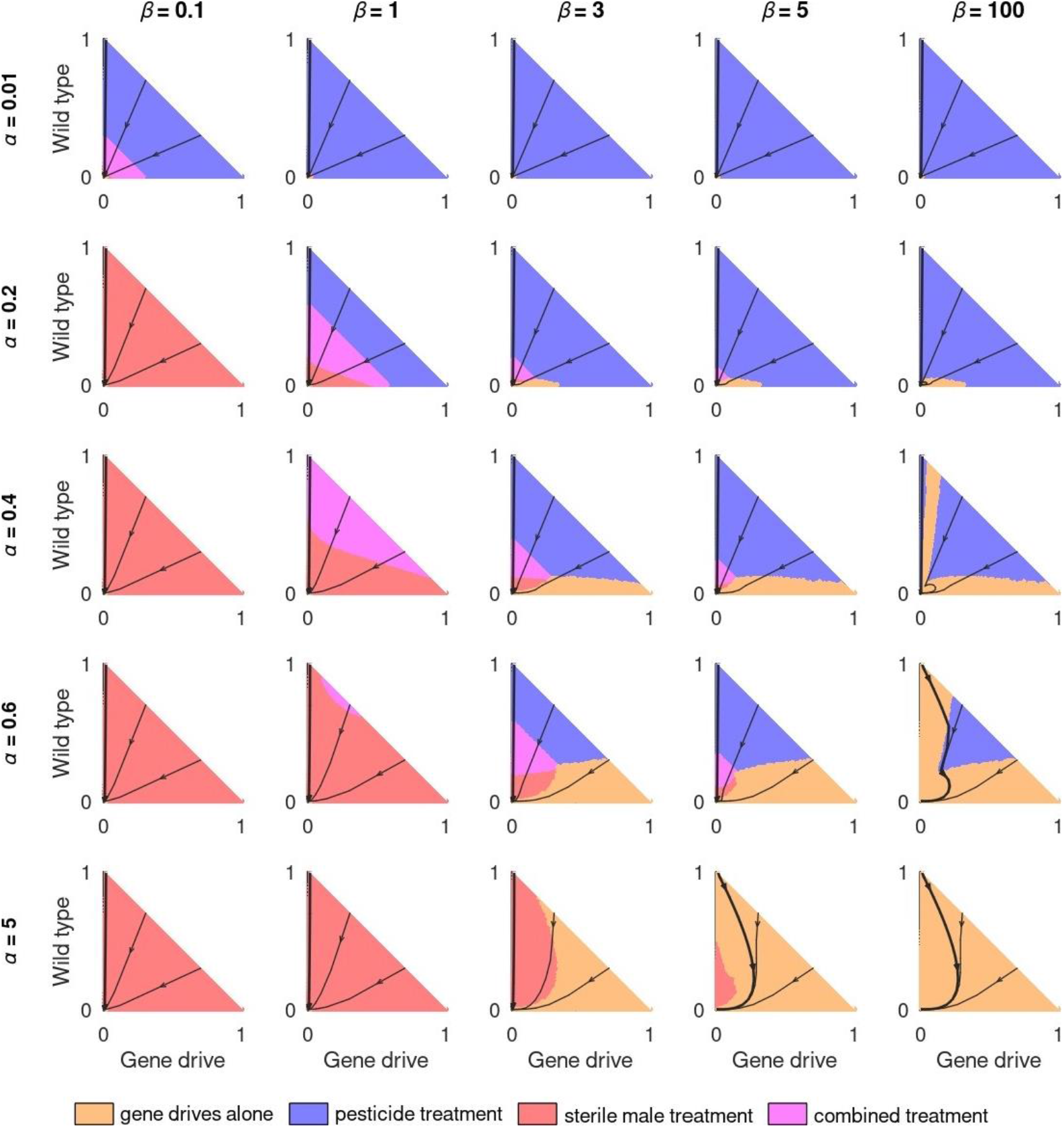
Population phase diagrams for cases in which both pesticides and sterile males can be used. Axes represent abundances of the two alleles: gene drive (x-axis) and wild-type (y-axis). The background colors denote states where gene drives are used alone (orange) or combined with additional treatments: pesticides (blue), sterile males (red), or both (magenta). The black curves show the population’s trajectories starting at different initial conditions (the bold curve corresponds to the standard deployment scheme). Parameters: *c* = 0.9, *s* = 0.7, *h* = 0.5 and *r* = 2.

When both methods are affordable enough to be considered, but not cheap enough to be used alone, the optimal treatment strategy typically consists of several stages. In that case, pesticides are applied to initially suppress the population. This occurs either immediately, together with the gene drive release, or after a few generations that are needed for the gene drive to reduce the mean growth rate of the population so that pesticides become effective. Later, sterile males are released to sustain the population at a low density until gene drives have spread sufficiently. From that point onward, gene drives complete the eradication process alone. This sequence is demonstrated in the trajectories passing from blue through magenta to red and ultimately anchoring in the orange area on the phase plane in Figure 5. This reflects the fact that pesticides are more effective in a large population, while sterile males are most effective in a small one (Lampert & Liebhold 2021).

### 3.4. Application of pesticides before gene drive deployment may provide an additional advantage

In the previous sections, we assumed that gene drives are released into a population that has reached its carrying capacity, and that treatments can be applied from that point onwards. Another possibility for using established pest controls is preliminary treatment—suppressing the wild-type population before introducing gene drives. Here, we examine this option for pesticide application, because sterile-male release is most cost-efficient in already suppressed populations (see Subsection 3.2). The intensity of preliminary pesticide application is subject to optimization due to an inherent trade-off: it facilitates eradication because gene drives start spreading from a higher initial frequency (as the population is already suppressed), but it requires additional investments. Notably, the timing of preliminary pesticide application does not require optimization, because it must always be most efficient to apply pesticides right before the gene drive release (earlier treatment would allow partial population recovery by the time the gene drives are released, and is therefore wasteful). Thus, here we optimize the intensities of possibly applying pesticides both prior and subsequent to the gene drive release, extending the results of Subsection 3.1.

Our results show that, when pesticides are relatively cheap (low values of *α*), the total cost is minimized when the preliminary application of pesticides is intense, and the population density after this preliminary application is low (Fig. 6A). In such cases, because the same amount of gene drive carriers is introduced to a relatively small population, gene drive frequency is initially high, leading to rapid eradication and low spillover risk. However, when pesticides are expensive, with values of *α* above a certain threshold, the optimal strategy implies no preliminary treatment (Fig. 6B).

**Figure 6.**
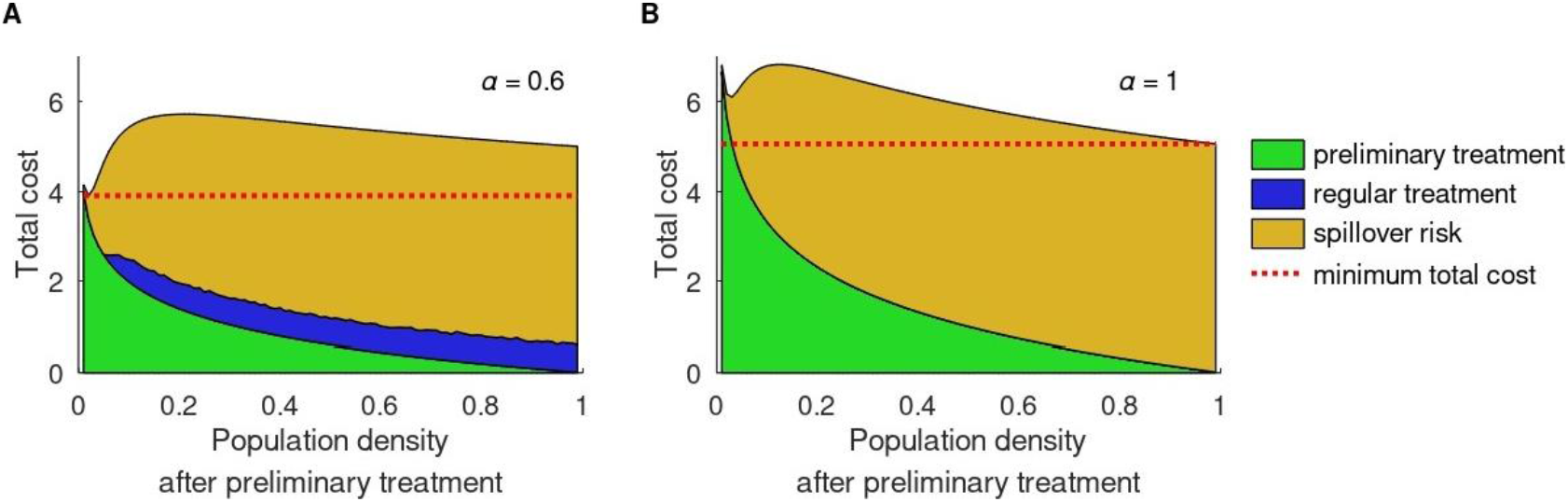
Optimal strategies for minimizing total cost of eradication when pesticides are applied both before and after gene-drive release. Shown are the total costs of eradication as a function of the population density after preliminary treatment, assuming optimal pesticide application from gene-drive release onward. The colors represent the components of the total cost: preliminary pesticide application (green), regular pesticide application (blue), and spillover risk (orange). Shown are two qualitatively different results, corresponding to the optimal solutions for different choices of *α* (dotted red lines): (A) Intense preliminary pesticide application is optimal, as seen in the total cost reaching its minimum at low population density (around 0.02); and (B) Having no preliminary treatment is optimal, as seen in the minimum cost occurring at population density 1 (carrying capacity). Parameters: *c* = 0.9, *s* = 0.7, *h* =0.5, *r* = 2, and initial condition is *n*_*D*,0_ = 0.01.

## 4. DISCUSSION

We developed a modeling framework for investigating the potential of integrating gene drives with established pest control methods to minimize spillover risk and the total cost of eradication. This framework allows incorporating gene drive deployment into multiple-treatment schemes, controlling for both direct expenditures and potential damage of spillovers. Our results show that both pesticide application and sterile-male release can substantially reduce spillover risk, and we demonstrated how the optimal treatment strategy depends on biological and economic parameters. Nevertheless, optimal use of treatments in combination with gene drives differs qualitatively between methods, with pesticides applied optimally early on and timed to coincide with maximal abundance of gene drive carriers, whereas sterile-male releases having two types of optimal application regimes, one intensive and one minimal.

Following all optimal strategies, regardless of the per-unit treatment costs, we observed a time point since which pesticide or sterile male interventions should cease, and gene drives finalize eradication alone (see Fig. 2A and Fig.3). This occurs because, when the gene drive is sufficiently frequent, the population density monotonically decreases, eventually leading to a reduction in gene drive carrier abundance. In turn, the population density is already low, implying low spillover, and therefore the marginal benefit of additional treatment becomes lower. This result aligns with previous findings in ecological restoration, where intervention occurs during intermediate stages and halts towards full recovery (Lampert & Hastings 2014, 2019). In both cases, interventions lead the system to a state from which the desired outcome, whether eradication or recovery, can be reached through system-internal dynamics alone, making additional treatments unwarranted. Additionally, we observed cases in which optimal strategies did not include established pest controls at the early stages of eradication. This occurs because, at early stages, when gene drives have only been introduced and have yet to substantially spread, spillover risk is still low, while the population growth rate is still high. Therefore, applying treatments during early stages would not substantially reduce the total spillover risk, and the population would be able to recover rapidly from the treatment.

Our study revealed an important distinction between pesticide applications and sterile-male releases in terms of their optimal use in combination with gene drives. With the increasing per-unit cost of pesticides, their total application gradually declined (see Fig. 2C). In contrast, a threshold value of the per-unit cost of sterile males emerged, which discriminates two types of strategies: (i) intensive releases of sterile males starting from the earliest stages (Fig. 3A, trajectories on the left) and (ii) limited, cheap releases at late stages of eradication (Fig. 3A, trajectories on the right). If both established pest controls complement gene drives, then pesticides should be applied first, when the population density is high, and sterile males released later, when the population density becomes lower (Fig. 5). Such temporal separation is also exhibited in the optimal combination of treatment methods without gene drives (Lampert & Liebhold 2021), and it occurs because pesticides are most cost-effective at higher population densities, whereas sterile males are most cost-effective at lower population densities, where their proportion is high enough to ensure predominant mating with sterile rather than wild males (Fister *et al*. 2013; Lampert & Liebhold 2021). The higher efficiency of sterile males at lower densities also implies that the entire eradication can be completed solely via sterile males, as demonstrated in cases where the per-unit cost of releasing sterile males is below the critical threshold (Fig. 3).

Our general approach was to identify optimal treatment strategies in terms of total costs of both treatment and spillover risk. At the same time, the posed problem can be viewed as a multiple-objective optimization problem, with spillover risk and expenditures on treatments being the two targets requiring simultaneous minimization. Assigning per-unit costs to these two targets and merging them into a single objective function—the total cost of eradication—would fall under the weighted-sum method of handling multiple-objective optimization. The total expenditure on treatment and the total spillover risk form the Pareto front, visualized as spillover-treatment curves in Fig. 2D and Fig. 3 (right panels). Such curves provide a useful practical tool because they eliminate the need to explicitly consider the per-unit treatment cost, which is often difficult to quantify. They demonstrate the best one could hope for: no solution exists that is both cheaper and results in lower spillover risk than any point on the Pareto front (Gunantara 2018). Note that the spillover-treatment curves (Pareto fronts) obtained for pesticides are continuous and convex, implying that any point on the Pareto front results from a unique value of the per-unit treatment cost, α (Fig. 2D). However, the spillover-treatment curves for sterile males contain separate segments, separated by dashed lined (Fig. 3, right panels), indicating regions that no not exhibit the optimal solution for any value of β. Specifically, in these regions, the Pareto front is concave, implying that no value in that segment could be optimal.

One of the challenges in formulating gene drive dynamics as an optimal control problem is that spillover risk is difficult to quantify. Therefore, our goal was not to assess optimal strategies with respect to an absolute level of spillover risk, but rather to evaluate the extent to which integrating established pest control methods can reduce spillover relative to not using additional treatments. Because we model spillover risk abstractly, we adopted a cautious approach that tends to favor overestimating rather than underestimating the risk. For this reason, we treat heterozygotes as posing the same level of risk as gene drive homozygotes (Eq. 7), despite the former carrying only one copy of the gene drive allele. Consistent with this precautionary approach, we focused on gene drives that inevitably spread through populations, even though threshold-dependent drives (Greenbaum *et al*. 2021; Leftwich *et al*. 2018), or other approaches of localization (Champer *et al*. 2020; Noble *et al*. 2019; Zhu & Champer 2023) present lower spillover risks.

Because optimal strategies combining gene drives with other controls have not been previously investigated, we employed a simplified model for tractability that does not incorporate stochasticity (Camm & Fournier-Level 2025; Li & Champer 2023), spatial structure (Beeton *et al*. 2022; Greenbaum *et al*. 2021; Harris & Greenbaum 2023; North *et al*. 2020), or type of selection (soft vs. hard; (Harris & Greenbaum 2023). Nevertheless, our modeling approach is flexible and can be extended to include other biological and ecological features, such as life cycles, spatial configurations, or environmental sensitivity (Kim *et al*. 2023). One important aspect that should be considered in future models is the possibility of resistance to gene drive evolving along the course of eradication. Resistance poses an additional risk in gene drive deployment that can be formulated as a cost, similar to how we treated spillover risk. Resistance risk may possibly interact with spillover risk in complex ways: on the one hand resistance may act as a natural safety mechanism, buffering spillover (Noble *et al*. 2018; Unckless *et al*. 2017), while on the other hand, it might result in stable polymorphism and persistence that can prolong spillover risk (Cook *et al*. 2022). Because the use of established pest controls may affect both suppression and resistance evolution, optimal strategies might utilize the advantages of different controls to minimize the different risks.

The integrated treatment schemes developed in this study provide an opportunity for advancing gene drive technology toward field trials and eventual full deployment by providing a framework for using established and well-known controls alongside gene drives. By providing a clearer understanding of how spillover risk can be reduced using available resources and treatment types, our framework supports more informed and responsible decision-making. This may help alleviate environmental damage and ease regulatory barriers that currently hinder the adoption of gene drives as a tool for addressing urgent public health challenges.

## Data Accessibility

The code used for simulations is publicly available at GitHub: https://github.com/sviatoslavrybnikov/optimal_treatment/

## Author Contributions

SRR conducted the analysis and wrote the manuscript. AL and GG conceived and supervised the study equally, with ordering of these authors determined via lottery. All authors commented and edited the manuscript.

## Acknowledgments

SRR was supported by the Minerva Center for the study of Population Fragmentation (MCPF). The study was supported by Israel Science Foundation (ISF) Grant 2049/21 awarded to GG and by Israel Science Foundation (ISF) Grant 1180/23 awarded to AL. We thank Keith D. Harris for his assistance with organizing and optimizing cluster computations, and Nataliya Rybnikova for her help in visualizing the study results.

## APPENDICES

## A. Derivation of equations for evolutionary and demographic dynamics

In this Appendix, we derive the recurrent equations for gene drive allele frequency, *q*_*t*_ (Eq. 1), and population density, *n*_*t*_ (Eq. 3). We denote the genotype abundances in the population at the beginning of generation *t* as *n*_*WW,t*_, *n*_*WDc,t*_, *n*_*WDnc,t*_, and *n*_*DD,t*_ representing the number of wild-type homozygotes, converted heterozygotes (i.e., heterozygotes for which the gene drive allele succeeded in conversion during gamete formation, so all produced gametes carry the gene drive allele), non-converted heterozygotes (i.e., heterozygotes for which gene-drive conversion during gamete formation failed), and gene drive homozygotes, respectively. At that moment, the frequency of the gene drive allele is, therefore,

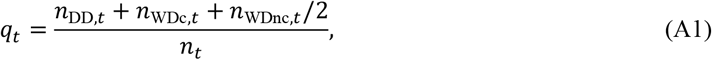

where *n*_*t*_ = *n*_*WW, t*_ + *n*_*WDc, t*_ + *n*_*WDnc, t*_ + *n*_*DD, t*_ is the total population size.

Following conversion and random mating, the numbers of the four genotypes change as follows:

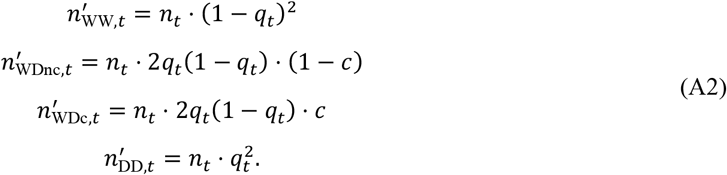

Assuming the Beverton-Holt growth model (Eq. 3), and adjusting it for reduced survival rates of gene drive carriers compared to the wild-type individuals (namely, 1 – *s* for gene drive homozygotes and 1 – *hs* for heterozygotes), we obtain the abundances of the four genotypes after selection, i.e. at the beginning of the next generation are:

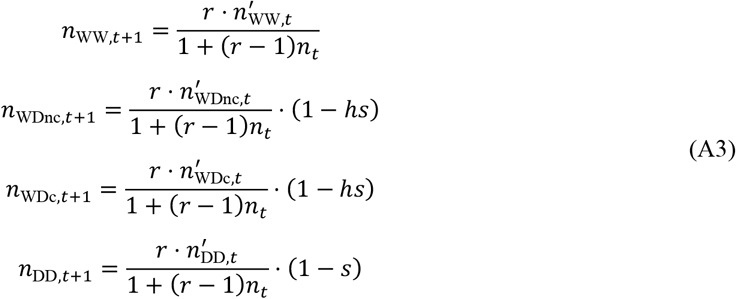

Placing genotype numbers from Eq. A3 into Eq. A1 gives the iterative formulae for the frequency of the gene drive allele:

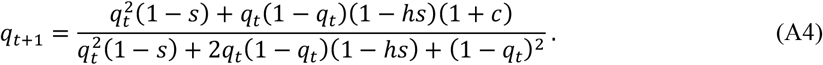

Summing up genotype numbers from Eq. A3 provides the total population size in the next generation:

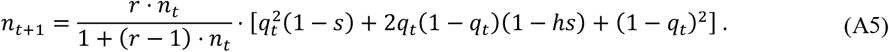

Since the denominator in Eq. A4 and the multiplier in Eq. A5 represent the population’s mean fitness, these equations can be written in the form of Eq. 1 and Eq. 3.

## B. Dynamic programming algorithm

In this Appendix, we describe the algorithm that we implemented and used to find the optimal solution. Specifically, we seek to find the amount of pesticide application in each generation, *P*_*t*_, and of sterile-male release in each generation, *M*_*t*_, that minimize the total cost of eradication over time (Eq. 8), considering the dynamics described in Eq. 1 and Eq. 4, 5, or 6, and the constraints *P*_*t*_ ≥ 0 and *M*_*t*_ ≥ 0 for all *t*. To find this optimal solution, we implemented a dynamic programming algorithm (Mangel & Clark 1988). The state variable is the population density, *n*_*t*_, represented by a finite set of discrete states, ranging from 0 to 1 with increments of 10^−4^. In turn, the gene drive allele frequency, *q*_*t*_, is not affected by the treatment used throughout the process: since *q*_*t*_ depends only on the frequency of gene drive carriers, not on *n*_*t*_ (Eq. 1), whereas treatment affects only *n*_*t*_ and not the frequency of gene drive carriers. Accordingly, we calculate *q*_*t*_ in advance and pass it to the algorithm as a pre-defined input function. To initialize the algorithm, we first estimate the duration of eradication achieved by gene drives alone, without any additional treatment. We round this duration up to the nearest ten, and use the resulting value *T* as the number of iterations in dynamic programming (Mangel & Clark 1988). For example, for the baseline gene drive configuration examined throughout the study, we obtained *T* = 40. This choice ensures that *T* is sufficiently large to guarantee the robustness of the algorithm. As a heuristic, we set the value function at *t* = *T* as minus the spillover cost (Eq. 7), 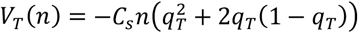.

Next, we use backward induction to find the value function, *V*_*t*_(*n*), and the optimal treatments, *P*_*t*_ and *M*_*t*_, for every state of the system, *n* = 0, 0.001, 0.002, … 1, and for all times, starting at *t* = *T* − 1, and going backward in time until *t* = 1. At each generation *t*, for each state *n*, we first use the gene-drive-adjusted Beverton-Holt model (Eq. 3) to find the population density *ñ*_*t*+1_ expected in the next generation under no treatment (rounded to the nearest state). Then, we examine the set of all possible interventions, from no treatment to complete eradication. For each state *n*_*t*+1_ resulting from these interventions (0 ≤ *n*_*t*+1_ ≤ *ñ*_*t*+1_), we select the intervention that maximizes the value function *V*_*t*_(*n*), defined as minus the sum of the three costs: (i) spillover cost, (ii) treatment cost, and (iii) the cost of the remaining treatment, from generation *t* + 1 onward, assuming the optimal strategy is applied from the next generation onward, which is given by −*V*_*t*+1_(*n*) and has been calculated at the former iteration of the algorithm.

Specifically, the spillover cost (i) is calculated from the gene drive allele frequency, *q*_*t*_, and population density, *n*_*t*_, according to Eq. 7. In turn, the treatment cost (ii) is calculated as follows. For treatment with pesticides, we use Eq. 4 to derive *P*_*t*_—the application required to suppress the population from *ñ*_*t*+1_ to *n*_*t*+1_. Specifically, it follows from Eq. 4 that 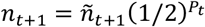, which implies

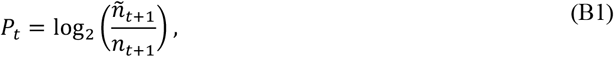

and the treatment cost is given by *C*_*P*_*P*_*t*_. Similarly, for treatment with sterile males, we use Eq. 5 to derive *M*_*t*_—the release required to move the population to the suppressed state *n*_*t*+1_ rather than to *ñ*_*t*+1_ expected under no treatment. Specifically, it follows from Eq. 5 that *n*_*t*+1_ = *ñ*_*t*+1_ · *n*_*t*_/(*n*_*t*_ + 2*M*_*t*_). Thus, we extract the term *n*_*t*_/(*n*_*t*_ + 2*M*_*t*_), which describes the effect of sterile males, and then solve for *M*_*t*_ from its denominator:

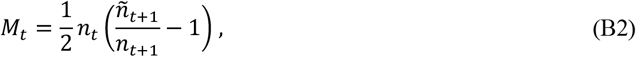

and the treatment cost is given by *C*_*M*_*M*_*t*_. Finally, for the combined treatment involving both methods, we can find the optimal pesticide application 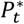 and the optimal sterile-male release 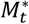 that together minimize the treatment cost, *C*_*P*_*P* + *C*_*M*_*M*, under the population dynamics constraint given by Eq. 6. The only non-trivial minimum of the treatment cost 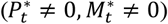 is given by:

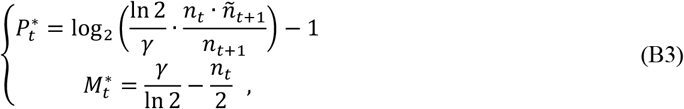

where *γ* = *α*/*β* = *C*_*P*_/*C*_*M*_. This non-trivial optimum satisfies both conditions *P*_*t*_ > 0 and *M*_*t*_ > 0 if and only if *γ* is within the range

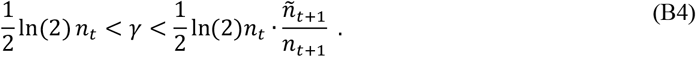

Otherwise, only one of the established pest controls should be used—either pesticides, if the ratio is below the bottom limit, or sterile males, if the ratio is above the upper limit; their magnitude is given as before by Eq. B1 (for pesticides) or Eq. B2 (for sterile males).

